# Structural and sequential regularities modulate phrase-rate neural tracking

**DOI:** 10.1101/2024.01.15.575585

**Authors:** Junyuan Zhao, Andrea E. Martin, Cas W. Coopmans

## Abstract

Electrophysiological brain activity has been shown to synchronize with the quasi-regular repetition of grammatical phrases in connected speech – so-called phrase-rate neural tracking. Current debate centers around whether this phenomenon is best explained in terms of the syntactic properties of phrases or in terms of syntax-external information, such as the sequential repetition of parts of speech. As these two factors were confounded in previous studies, much of the literature is compatible with both accounts. Here, we used electroencephalography (EEG) to determine if and when the brain is sensitive to both types of information. Twenty native speakers of Mandarin Chinese listened to isochronously presented streams of monosyllabic words, which contained either grammatical two-word phrases (e.g., catch fish, sell house) or non-grammatical word combinations (e.g., full lend, bread far). Within the grammatical conditions, we varied two structural factors: the position of the head of each phrase and the type of attachment. Within the non-grammatical conditions, we varied the consistency with which parts of speech were repeated. Tracking was quantified through evoked power and inter-trial phase coherence, both derived from the frequency-domain representation of EEG responses. As expected, neural tracking at the phrase rate was stronger in grammatical sequences than in non-grammatical sequences without syntactic structure. Moreover, it was modulated by both attachment type and head position, revealing the structure-sensitivity of phrase-rate tracking. We additionally found that the brain tracks the repetition of parts of speech in non-grammatical sequences. These data provide an integrative perspective on the current debate about neural tracking effects, revealing that the brain utilizes regularities computed over multiple levels of linguistic representation in guiding rhythmic computation.

## 1. Introduction

Language comprehension requires the transformation of a continuous physical signal (speech or sign) into discrete structured representations (viz., linguistic units and structured relations between them). Evidence shows that during successful comprehension, low-frequency neural activity becomes aligned with linguistic units such as phonemes and syllables^1,2^. This “neural tracking” effect was initially shown for these lower-level linguistic units, but more recently it has been observed for higher-level and rather abstract units such as words and phrases as well [3–9]. While syllables can be physically characterized by salient energy increases^10^, these higher-level units have no one-to-one physical correlates in the speech signal, meaning that they require endogenous computations to support neural tracking^11,12^.

It has been established that the brain tracks both *sequential* and *structural* linguistic information in spoken language. That is, when a connected speech stream contains a regular sequential pattern of (certain subtypes of) words or syllables, the brain response will show neural tracking at the rate at which these linguistic units occur in the speech^5,13,14^. What is striking is that structurally organized information elicits similar effects. In particular, when phrasal units like verb phrases or noun phrases are regularly repeated in a connected speech stream, the brain response will show neural tracking at the phrase rate^3,6–9^. Since phrase structure is organized hierarchically, these structural regularities are not directly visible in the sequential structure of language – the notion ‘phrase rate’ presumes the cognitive capacity to represent hierarchical phrase structure by abstracting away from the sequential input. It thus appears that the brain concurrently tracks linguistic information at multiple levels of representation (i.e., syllables, words, phrases) and at multiple timescales^5^. Throughout our discussion of neural tracking, we will use the terms ‘structural information’ and ‘sequential information’ to denote the contrast between linguistic information that is hierarchically organized (in particular, compositional phrase structures) and linguistic information that is sequentially organized in speech (in particular, the sequential repetition of lexical information). While we note that the contrast is not dichotomous, we keep this description because it captures one of the fundamental challenges of language processing: as hierarchical structure is not directly encoded in the linear speech input, tracking hierarchical structure requires processes of perceptual inference that go beyond merely following the sequential input^11,15^.

Because sequential distribution is (partially) determined by structural properties, structural and sequential regularities in natural language are correlated. For instance, syntactic structure has correlates in the surface statistics of sentences^16–18^, and processing syntactic structure can give rise to statistical expectations about sequential properties of sentences (e.g., the likelihood that certain words are adjacent^19,20^). Due to the correlation between structural and sequential information, it is difficult to delineate their functional roles in language processing, in particular when viewed through the lens of neural speech tracking^9,21^. In this electroencephalogram (EEG) study, we address the role of sequential and structural information in neural speech tracking by presenting participants with speech streams that vary in syntactic structure. In particular, we ask (1) whether structural and sequential regularities affect tracking differentially, and (2) how syntactic variables modulate the neural tracking of phrases.

### 1.1 Neural speech tracking is modulated by structural information

Because natural language is hierarchically structured, it is rife with properties that separate meaning from the linear or temporal order in which it is produced^22–26^. Processing language therefore requires internal computations that integrate and resolve information over temporal displacement and variation^11,15^. Such internal computations allow basic elements to break free from the linear order they are imposed with and be interpreted with respect to hierarchically related units. In this section, we will discuss a growing body of research that shows a connection between hierarchical structure and the neural tracking of speech.

The first attempts to reveal the role of hierarchical structure in neural tracking are attributed to Ding and colleagues^4,5^. In their “frequency-tagging” studies, participants listened to sequences of monosyllabic words, which were isochronously presented at 4 Hz. In these sequences, two adjacent words could be grouped into two-word phrases (occurring at 2 Hz), and two adjacent phrases could be grouped into four-word sentences (occurring at 1 Hz). As the sequences contained synthesized speech that was presented without prosody or coarticulation, neither phrases nor sentences were auditorily cued in the stimulus, and their rates were frequency-tagged only by virtue of participants’ knowledge of grammar. The team found that electrophysiological brain activity closely matched the occurrence rates of these abstract linguistic units: phrases elicited spectral peaks at 2 Hz, and sentences elicited a spectral peak at 1 Hz (see also other evidence^27,28^). Crucially, this frequency-tagged effect of grammatical structure disappeared when participants listened to structurally identical materials in an unfamiliar language. These findings suggest that structural knowledge of language supports “chunking” continuous speech into structured, multi-word units^29^.

In addition to such abstract grouping patterns, further studies showed that the content of linguistic structure also modulates neural speech tracking. Using naturally produced speech, two studies^6,7^ showed that phrases were tracked more strongly in sentences than in prosodically natural jabberwocky sentences and in unstructured word lists. Moreover, studies have found that the strength of phrase-rate tracking is correlated with behavioral measures of comprehension^30,31^, thus showing the perceptual relevance of neural tracking. Note that by phrase-rate tracking we mean tracking effects at the frequency that corresponds to the occurrence rate of phrasal units (i.e., at 1 Hz in our experiment). It can, but need not, refer to the tracking of syntactic phrases.

That neural tracking of speech is indicative of a structural specification extending beyond the contrast between sentences and word lists is also suggested by a recent frequency-tagging study. A recent study^3^ reported stronger tracking at the phrase rate when participants listened to sequences of structurally consistent adjective-noun phrases than when they listened to sequences that were composed of structurally-variable two-word phrases (e.g., adverb-adjective, verb-noun, preposition-noun, etc.). Their findings are interesting in two respects. First, the fact that phrases were tracked even when they were composed of different parts of speech shows that a purely sequential account of phrase-rate tracking is insufficient, because these stimuli contained no sequential regularities at the phrase rate. Instead, the only phrase-rate regularity in these variable sequences is a structural one, namely the repetition of two-word phrase structures. Second, the fact that phrases in structurally variable sequences were nevertheless tracked less closely than phrases in structurally consistent sequences shows that, even in the context of intact syntactic composition, the structural details of a phrase influence phrasal tracking. However, as the structurally-variable condition in this study^3^ differed from the structurally-consistent adjective-noun condition in various linguistic respects (e.g., type of attachment, position of the head, see Section 1.3), it remains unclear which linguistic factors modulate phrasal tracking. As studies that explicitly examine the role of syntactic or semantic information are sparse, a gap remains between our rather coarse-grained understanding of neural speech tracking and our fine-grained knowledge about linguistic structure^32^.

### 1.2 Neural speech tracking is modulated by sequential information

Identified as one confound in the original grammatical tracking study^5^, the regular, sequential occurrence of lexical information has been argued to be a more parsimonious explanation of phrase-rate tracking^13,14^. According to this view, a tracking effect emerges at the phrase rate because certain word-level features repeat in sequences at the same frequency as phrases do. A 2-Hz phrase-rate peak arises when participants listen to isochronously presented 1-second sentences like “dry fur rubs skin” simply because nouns (i.e., “fur”, “skin”) are repeated every 1/2 second^13^. This account of tracking is based on the sequential occurrence of word-level information (i.e., the semantic features associated with parts of speech, derived from statistical distributions) and therefore does not require hierarchically structured representations.

Another non-hierarchical account of phrase-rate tracking holds that the effect is driven by sequential statistics, that is, the rhythmic temporal variation in transitional probabilities between words. In the sentence “dry fur rubs skin”, word-to-word transitional probabilities are higher within the phrases “dry fur” and “rubs skin” than between these two phrases (i.e., at the boundary between “fur” and “rubs”), which leads to an alternating high-low pattern that coincides with phrase occurrence (cf.^18^). Given that the human brain is sensitive to transitional probabilities during speech segmentation (to extract word-like units^33^), it might employ similar mechanisms to extract phrases in speech sequences. Indeed, similar tracking effects are found when participants listen to connected sequences of unfamiliar words that are cued only by high transitional probabilities between syllables^34–36^. For example, in one study^34^ Dutch native speakers were presented with sequences of disyllabic Chinese words. As these participants had no prior knowledge of Chinese, the only cues available to extract multi-syllabic units were the differences in syllable-to-syllable transitional probabilities, which were high within words and low between words. Over the course of the experiment, tracking at the word rate emerged for these statistically indexed words. In order to show that it is linguistic structure that drives neural tracking in a frequency-tagging design, it is therefore important to control for the differences between conditions in terms of the sequential transitional probability between linguistic units.

### 1.3 The present study

The previous two sections show that both structural and sequential regularities correlate with rhythmic neural activation. In the present study, we seek to enrich our understanding of these relationships by presenting participants with speech streams that vary in syntactic structure and sequential-statistical information. By measuring phrase-rate tracking in these different speech streams, we investigate (1) how structural (syntactic) and sequential regularities together affect the neural tracking of speech, and (2) which aspects of syntactic structure are reflected in tracking effects. To answer the first question, we used a balanced design in which structural and sequential information were orthogonalized as much as possible. We say as much as possible because a full orthogonalization between structural and sequential information can hardly be achieved since these variables are highly correlated in natural language. Specifically, half our conditions contain grammatical word combinations, the other half contain non-grammatical combinations. Moreover, half of our conditions contain a sequential regularity at the phrase rate (i.e., the sequential repetition of lexical information), the other half do not. The structural and sequential factors were fully crossed, such that our design contained all combinations of +/- syntactic structure and +/- lexical repetition.

In the three grammatical conditions, we probed the effects of structural information neural tracking of phrases. The grammatical conditions consisted of repetitions of two-word phrases, in which we varied two properties that play a prominent role in syntactic theory. In particular, we varied the position of the head of the two-word phrase (Head position) and the syntactic function of the word attached to the head (Attachment type).

Syntactic structures are endocentric, which means that the syntactic status of a phrase is determined by one of its elements, the ‘head’. Thus, the verb is the head of a verb phrase (VP; e.g., *win* in *win the game*), the preposition is the head of a prepositional phrase (PP; e.g., *on* in *on Monday*), and so on. Moreover, the syntactic function of the other elements in a phrase is dependent on the selectional properties of the head. Elements that are required to complete the meaning of the head and receive a thematic role are called complements (or arguments). Adjuncts, instead, do not receive a thematic role; they modify the grammatical head and typically convey optional information^37,38^. Consider the VP *win the game on Monday.* The head of the phrase, the verb *win*, assigns a thematic role to *the game*, which is the verb’s complement (*the game* completes the meaning of *win*). The PP *on Monday* is not inherently required by the verb and does not receive a thematic role, so it is considered an adjunct. Certain syntactic theories additionally propose a difference between complement and adjunct attachments in terms of their underlying syntactic structure^38,39^.

Typologically, languages differ in whether the head occupies the left or right edge of the phrase. In a head-initial language like English, the head consistently precedes its complements. Adjuncts, on the other hand, are freer and can often be placed on either side of the head. These syntactic differences in attachment and head position and are visually illustrated in Figure 1.

**Figure 1.**
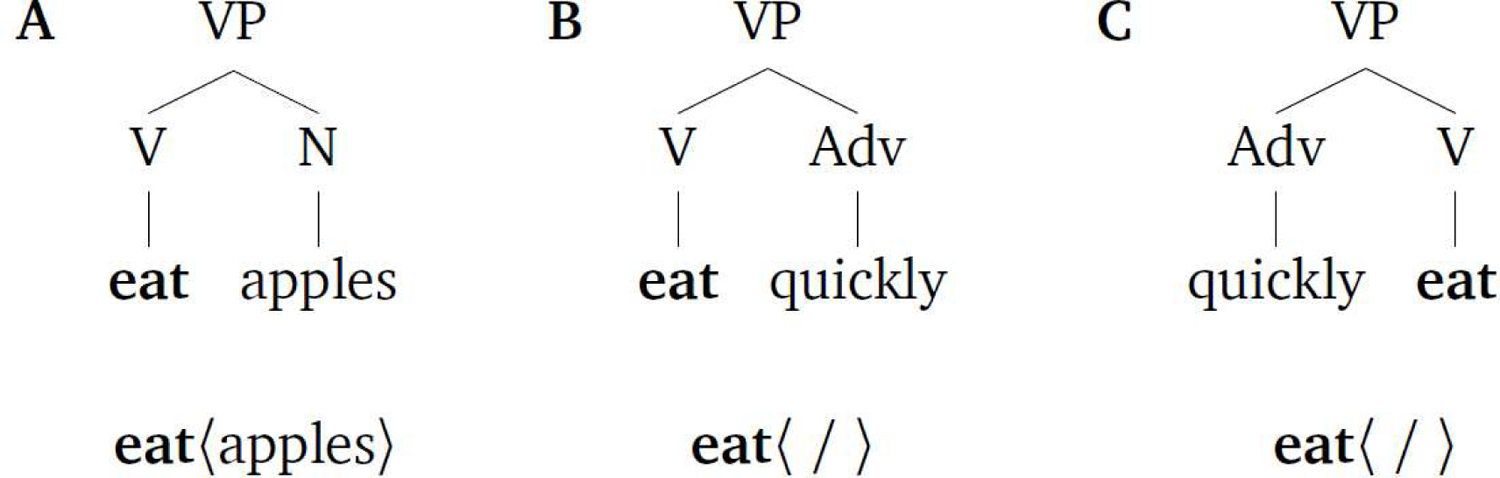
Syntactic representations of different verb phrases. In **(A)**, the noun *apples* is the complement of the verb *eat*, which is the head of the VP (in bold). In **(B)**, the adverb *quickly* is attached to the verb phrase as an adjunct. Because the phrase contains no complement, the thematic role of the verb is not assigned, represented as 〈 / 〉. The verb phrase in **(C)** is the same as the one in (B) except that the adverb precedes the verb, which appears in the final position.

We included the factors Attachment type and Head position for two reasons: **First**, the structurally variable ‘mixed-phrase’ condition in Burroughs et al. (2021) contained phrases that varied in these respects. Recall that they found stronger phrase-rate tracking in sequences containing structurally consistent adjective-noun (A-N) phrases than in sequences containing mixed phrases. An example of their mixed-phrase condition is given below:

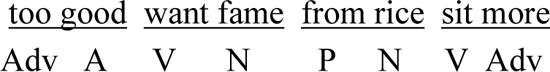

The position of the head of these two-word phrases is not consistent: the adjective *good* in the adjectival phrase (AP) *too good* is the second word, the verb *want* in the VP *want fame* is the first word, the preposition *from* in the PP *from rice* is the first word, and so on. Moreover, the word attached to the head in each phrase does not consistently fulfill the same syntactic function; it can be a complement or an adjunct. The adverb *too* is an adjunct to *good*, *fame* is the complement of *want*, *rice* is the complement of *from*, etcetera. Inconsistency in either of these variables (Head position and Attachment type) might have led mixed phrases to be tracked less closely than structurally consistent A-N phrases. We consider this a reasonable possibility because both variables affect language processing.

**Second**, both attachment type and head position have processing correlates. In head-initial structures, complements are easier to process than adjuncts. For instance, the PP *for a raise* is read faster than the PP *for a month* when preceded by *The company lawyers considered employee demands…*^40–44^. The noun *demands* takes a PP complement. Reading *demands* therefore facilitates the processing of *for a raise*, which can fulfill that complement role (cf. *to demand a raise*), compared to *for a month*, which is a temporal adjunct to the verb *considered*. This example also illustrates the role of the head (here, *demands*) in syntactic processing, which allows syntactic structure to be projected (as in head-driven parsing models^40,45^) and can trigger expectations for upcoming linguistic material^41,44,46^. Moreover, cross-linguistic differences in head position modulate sentence-processing effects. In head-initial structures it is commonly found that processing costs are increased when the distance between the head and its dependents is increased^47,48^. Studies investigating head-final structures, instead, have found the exact opposite, i.e., a facilitation in processing of words integrating longer dependencies, which is typically attributed to an increase in the prediction for the upcoming head^49,50^. Given the robust effects of Attachment type and Head position on sentence processing, we incorporated both factors in our design.

Constructing minimal contrasts with these two factors resulted in three grammatical conditions, whose labels refer to the parts of speech they are composed of: V-N, V-Adv and V-Adv-Alt. The V-N condition contains sequences of Verb-Noun combinations, which are consistently head-initial and contain a noun as the complement of the verb (like in Figure 1A). V-Adv sequences contain Verb-Adverb phrases that are consistently head-initial as well, but the adverb is an adjunct to the verb phrase (like in Figure 1B). In a third condition, V-Adv-Alt, sequences are composed of Verb-Adverb (adjunct) combinations whose order was shuffled, creating a random alternation between head-initial and head-final phrases (Figures 1A and 1C, respectively). Our stimuli were in Mandarin Chinese because it naturally allows phrases of both the V-Adv and the Adv-V order. In sum, the effect of Attachment type (complement vs. adjunct) is tested in the comparison between V-N and V-Adv. The effect of Head position (consistent vs. varying head position) is tested in the comparison between V-Adv and V-Adv-Alt.

In three non-grammatical conditions, we probed the effects of non-structural information, i.e., the sequential repetition of lexical information. The N-R condition contains sequences composed of two-word pairs with a Noun and a Random word, yielding sequential repetition of the noun at the phrasal rate in the absence of phrasal structure. The V-V consists of random Verb sequences, thus having sequential repetition of verbs at the word rate. The R-R condition contains random word sequences and therefore contains no regular repetition of lexical information at all. All non-grammatical conditions were constructed such that there were no grammatical phrases in the concatenated sequence. An overview of the experimental design with example materials is given in Table 1.

**Table 1.** Overview of the experimental design, with examples of each condition. The three lines in the Examples column represent Chinese characters, Chinese phonological transcription with tones, and word-by-word translation in English, respectively. In the grammatical conditions, the head of each verb phrase is indicated in bold.

| Condition | Structural regularity | Syntactic specification | Sequential regularity | Examples |  |
| --- | --- | --- | --- | --- | --- |
| V-N | ✓ | Head-Initial Complement | ✓ | 看天<br>kan4 tian1<br>**look** sky | 喂鱼<br>wei4 yu2<br>**feed** fish |
| V-Adv | ✓ | Head-Initial Adjunct | ✓ | 放下<br>fang4 xia4<br>**put** downwards | 派出<br>pai4 chu1<br>**send** outwards |
| V-Adv-Alt | ✓ | Head-random Adjunct | ✗ | 重做<br>chong2 zuo4<br>again **do** | 说明<br>shuo1 ming2<br>**state** clearly |
| N-R | ✗ | - | ✓ | 电伪<br>dian4 wei3<br>electricity fake | 发再<br>fa4 zai4<br>hair again |
| R-R | ✗ | - | ✗ | 下讲<br>xia4 jiang3<br>down talk | 变降<br>bian4 jiang4<br>change descend |
| V-V | ✗ | - | ✗ | 取听<br>qu3 ting1<br>get listen | 剥搬<br>bo1 ban1<br>peel move |

We recorded the EEG of participants while they listened to Mandarin Chinese speech streams that vary in structural and sequential regularities. The streams consisted of connected syllables isochronously presented at 2 Hz, yielding a 1-Hz rate for two-word phrases. Following previous work^3,4,8^, neural tracking was quantified through measures of evoked power and inter-trial phase coherence (ITPC) of the frequency-domain representation of the EEG signal.

We consider three theoretical possibilities, which make different predictions about the presence of a tracking effect at the 1-Hz phrase rate. First, a *structure*-only hypothesis posits that the brain only tracks syntactic structure. Following this hypothesis, a phrase-rate tracking effect is expected only in the conditions that contain syntactic structure (i.e., the grammatical conditions V-N, V-Adv, and V-Adv-Alt). We also hypothesize that our structural manipulations will yield different magnitudes of tracking^3,6^. However, we cannot make specific predictions about the directions of these effects, because in existing data several factors co-occur (e.g., in Burroughs et al., 2021, the adjective-noun and mixed-phrase conditions differed in multiple respects), and because mechanistic linking hypotheses between frequency representations and syntactic structure building are missing^32^. Second, a *sequence*-only hypothesis posits that the brain only tracks the sequential repetition of lexical information. It therefore predicts phrase-rate tracking effects in conditions with regular repetition at the word level (i.e., V-N, V-Adv, and N-R), regardless of whether these conditions contain syntactic structure as well. The *structure*-only and the *sequence*-only hypotheses may appear too simplistic, because it is unlikely that the brain only tracks one type of information. We nevertheless present the hypotheses this way because existing accounts of neural tracking effects lean towards one of the two explanations^8,13,14^. A third, *hybrid* hypothesis holds that the brain tracks both structural and sequential regularities. It predicts phrase-rate tracking effects in all conditions that contain regularity at the phrase rate, whether its source is structural or sequential (i.e., the grammatical conditions V-N, V-Adv, V-Adv-Alt, and the non-grammatical condition N-R), but does not specify the exact source of a tracking effect in cases where both types of regularity co-exist (i.e., in V-N and V-Adv). We consider the hybrid hypothesis most likely *a priori*, but nonetheless aim to assess the degree to which these three competing hypotheses are supported by our results. Figure 2 presents the predicted effects following the three hypotheses. As can be seen, the three hypotheses converge for conditions V-N, V-Adv, R-R, and V-V regarding the presence or absence of tracking effects. Given that they diverge only for V-Adv-Alt and N-R, these conditions will be the main focus of this study, in accordance with the research questions outlined above.

**Figure 2.**
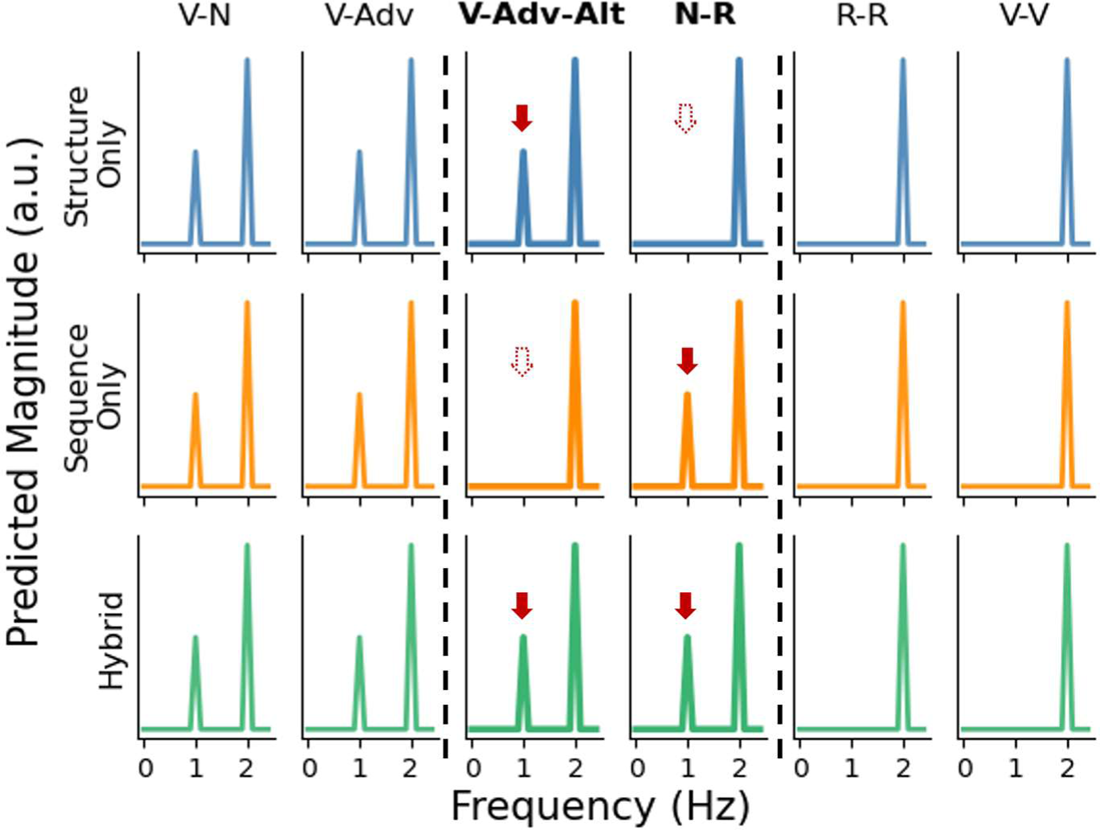
Predicted outcomes in the frequency domain following the three hypotheses. The predictions only indicate the presence or absence of peaks and do not reflect effect sizes. The columns for V-Adv-Alt and N-R are emphasized because these are the critical conditions for which the three accounts make different predictions. The solid red arrows in these two columns indicate phrasal peaks that are expected under the different hypotheses. Dashed arrows indicate the absence of spectral peaks at the phrasal frequency.

## 2. Results

At the 2-Hz syllable rate, all conditions elicited peaks both power and ITPC (all p < 0.001). A further comparison between conditions did not show any reliable difference in power or ITPC of the 2-Hz syllable peaks (all p > 0.4). Figures 3A and 3C presents the power and ITPC spectra of all conditions.

**Figure 3.**
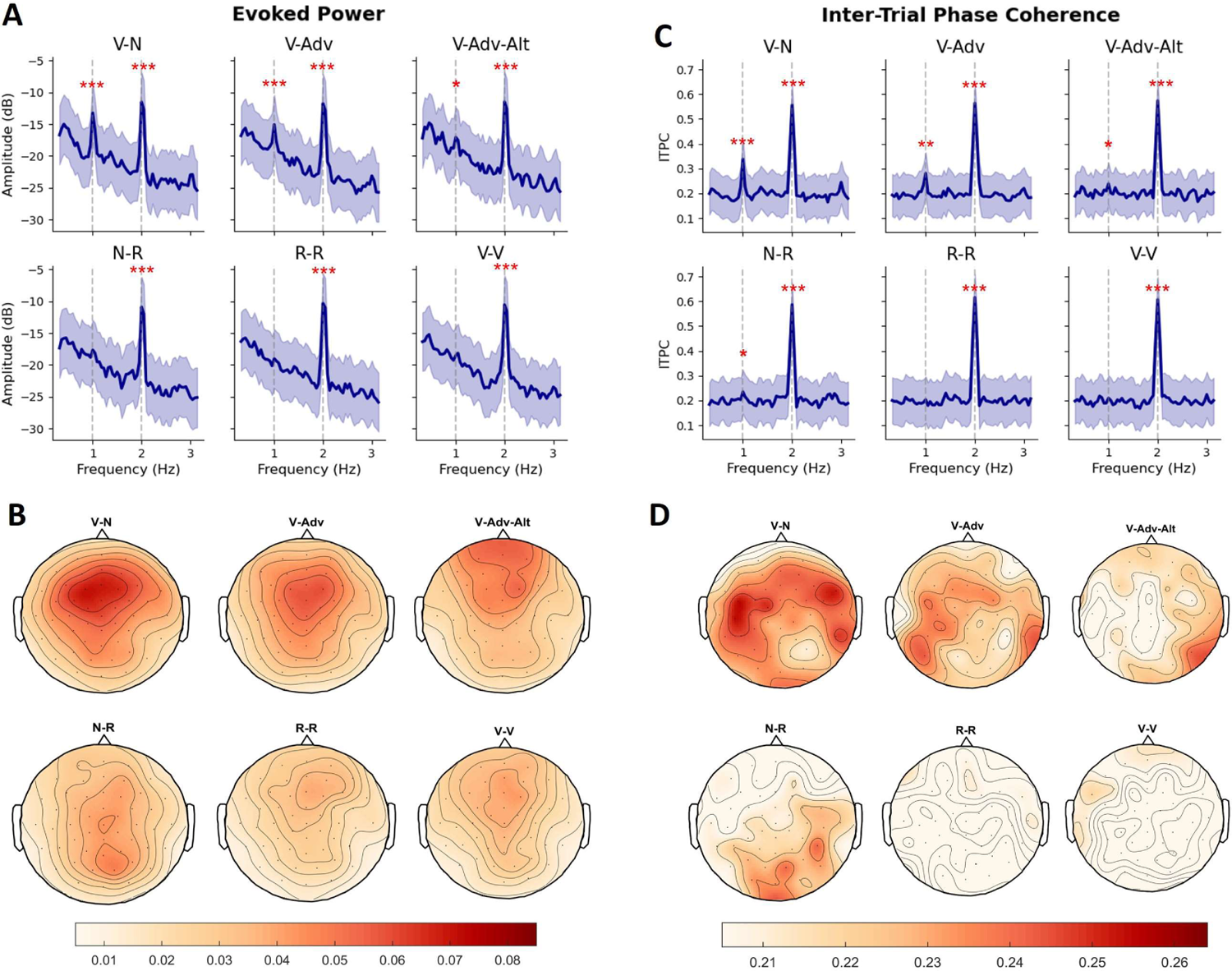
EEG responses in all conditions. **(A)** Spectra of evoked power. **(B)** Topography of 1-Hz evoked power. **(C)** Spectra of ITPC. **(D)** Topography of 1-Hz ITPC. In (A) and (C), the shaded areas indicate one standard error around the mean over participants. Significance: *, p < 0.05; **, p < 0.01; ***, p < 0.001.

At the 1-Hz phrase rate, the results for evoked power and ITPC were slightly different. For evoked power (Figure 3A), significant phrase-rate peaks were present in all grammatical conditions (V-N: p < 0.001; V-Adv: p < 0.001; V-Adv-Alt: p = 0.047). We did not find a phrasal peak in the power spectrum of the N-R condition (p = 0.46). For ITPC (Figure 3C), phrase-rate peaks were again found in the grammatical conditions V-N (p < 0.001), V-Adv (p = 0.002), V-Adv-Alt (p = 0.044), but also in the non-grammatical N-R condition (p = 0.043). Besides this difference in frequency spectra, power and ITPC at 1 Hz also had different scalp distributions (see Figures 3B and 3D). Differences between power and ITPC are not entirely unexpected, since ITPC has been shown mathematically to be more sensitive to neural responses that are synchronized to external stimuli^51^. This, combined with the fact that four out of six conditions showed a 1-Hz peak in ITPC (compared to three out of six conditions showing a 1-Hz peak in evoked power), reveals the higher sensitivity of ITPC to tracking effects. We will therefore focus on the pairwise comparisons of ITPC.

To examine whether neural tracking at the phrase rate was different across conditions, we performed pairwise paired samples t-tests on all conditions that showed a 1-Hz peak in the ITPC spectrum (i.e., V-N, V-Adv, V-Adv-Alt, and N-R). Based on two theoretically informed planned contrasts (i.e., V-N vs. V-Adv for the effect of Attachment and V-Adv vs. V-Adv-Alt for the effect of Head position), we found that phrases with complement attachments were tracked more strongly than phrases with adjunct attachments (V-N > V-Adv, Δ = 0.057, p = 0.034, see Figure 4A). Moreover, phrases with a consistent head position were tracked more strongly than phrases with a varying head position (V-Adv > V-Adv-Alt, Δ = 0.042, p = 0.012). We also found stronger tracking for V-N than for both V-Adv-Alt (Δ = 0.010, p < 0.001) and N-R (Δ = 0.103, p < 0.001), and stronger tracking for V-Adv than for N-R (Δ = 0.046, p = 0.008). There was no difference between V-Adv-Alt and N-R (Δ = 0.004, p = 0.772).

**Figure 4.**
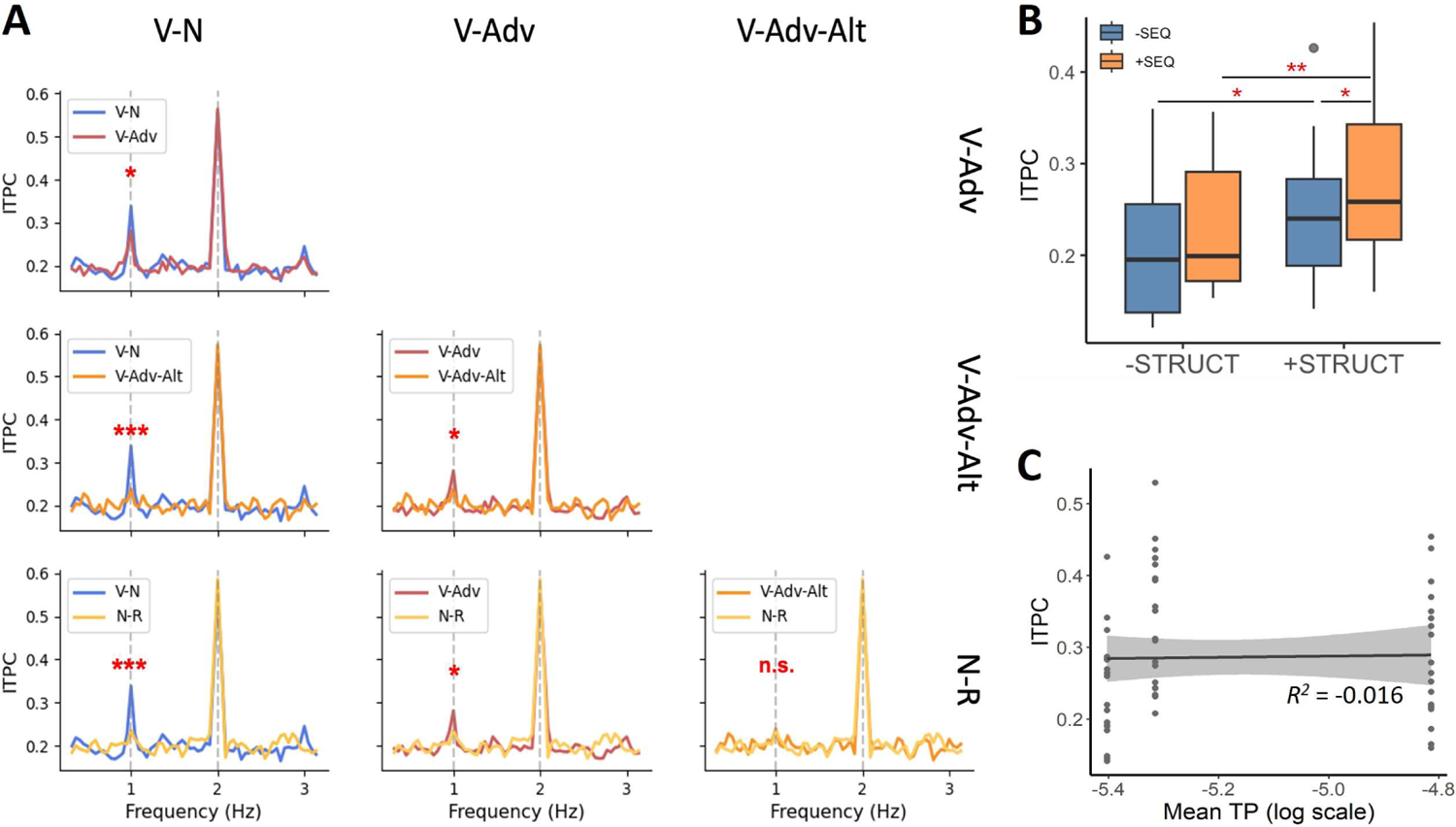
**(A)** Pairwise comparisons of 1-Hz peaks in ITPC. **(B)** Boxplot of ITPC grouped by STRUCTURAL and SEQUENTIAL regularities, where [+SEQ, +STRUCT] = V-Adv, [-SEQ, +STRUCT] = V-Adv-Alt, [+SEQ, -STRUCT] = N-R, and [-SEQ, -STRUCT] = R-R. Asterisks indicate the results of post-hoc comparisons following a two-way repeated measures ANOVA. **(C)** Scatter plot and least-squares linear regression fitted between ITPC and TP in all grammatical conditions. Mean TP values of each condition were used as the predictor. The shaded area indicates 95% confidence interval. Significance: *, p < .05; **, p < .01; ***, p < .001.

The ITPC results show that the brain is sensitive to both sequential and structural regularities. The next question is whether these effects are independently additive or whether they interact with one another. To test this, we ran a two-way repeated measures analysis of variance (ANOVA) on a subset of the data (see Figure 4B). In this subset, the two factors STRUCTURAL and SEQUENTIAL regularities were fully orthogonalized, yielding the following four groups: V-Adv ([+SEQ, +STRUCT], V-Adv-Alt [-SEQ, +STRUCT], N-R [+SEQ, -STRUCT], and R-R [-SEQ, - STRUCT]). As expected, the ANOVA yielded significant main effects of both structural (*FSTRUCT* (1, 19) = 13.13, *p* = 0.002) and sequential regularities (*FSEQ* (1, 19) = 6.86, *p* = 0.017). While, the interaction was not significant (*Finteraction*(1, 19) = 0.33, *p* = 0.572), the results in Figure 4B nevertheless suggest that the effect of SEQUENTIAL regularity is not the same for conditions with vs. without STRUCTURAL regularity. A post-hoc analysis on estimated marginal means^52^ tentatively confirmed this suspicion. Regarding the role of sequential regularity, when the stimuli contained grammatical structure, adding sequential regularity increased ITPC ([+SEQ, +STRUCT] > [-SEQ, +STRUCT], *β* = 0.042, *SE* = 0.014, *t*(19) = 2.85, *p* = 0.012). However, this effect of sequential regularity vanished when the stimuli contained no grammatical structure ([+SEQ, -STRUCT] = [-SEQ, -STRUCT], *β* = 0.032, *SE* = 0.019, *t*(19) = 1.68, *p* = 0.140). The role of structural regularity, instead, appears to be more consistent. Adding structural regularity increased ITPC, both when there was sequential regularity in the stimuli ([+SEQ, +STRUCT] > [+SEQ, -STRUCT], *β* = 0.046, *SE* = 0.016, *t*(19) = 2.96, *p* = 0.008) and when there was no sequential regularity in the stimuli ([-SEQ, +STRUCT] > [-SEQ, -STRUCT], *β* = 0.036, *SE* = 0.014, *t*(19) = 2.61, *p* = 0.017). All *p*-values reported for the post-hoc analysis were adjusted with Tukey’s method. Thus, it appears that the effect of STRUCTURAL regularity was more robust than the effect of SEQUENTIAL regularity: the effect of structural regularity showed up even in the face of sequential inconsistency, but the effect of sequential regularity was dependent on the presence of structure. We do note that these post-hoc pairwise comparisons should be interpreted with caution, because the interaction in the two-way ANOVA was not significant.

As an additional control analysis, we used a linear regression analysis to test if the 1-Hz ITPC effects can be explained by the transitional probabilities (TPs) of the two-word combinations in the different conditions. This analysis included the conditions V-N, V-Adv, and V-Adv-Alt. The condition N-R was not included because there were not enough N-R bigrams in the Wikipedia corpus to compute a sufficiently representative average TP for these non-grammatical combinations. The bigram TP of a phrase *w1w2* was computed as the probability of encountering that phrase divided by the probability of encountering all other two-word phrases starting with *w1*, or 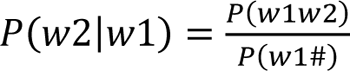. Word frequencies were extracted from the P(w1#) Chinese Wikipedia corpus used in a previous study^13^. ITPC on individual trials and average TP in the V-N, V-Adv, and V-Adv-Alt conditions were then entered into a linear regression model, which showed that TP did not reliably predict ITPC (Figure 4C; *R*^2^ = −0.016, *β* = 0.22, *SE* = 2.3, *t*(58) = 0.097, *p* = 0.923). Thus, while there might be differences between the conditions in terms of sequential statistics, this potential confound cannot explain the pattern of results we observed.

## 3. Discussion

In this EEG study, we investigated whether and how neural tracking of phrases in connected speech is modulated by regularities stemming from syntactic structure and from sequential lexical information. Besides robust syllable tracking in all conditions (Giraud & Poeppel, 2012), we observed tracking effects at the phrase rate in all conditions that contained a structural and/or sequential regularity at that rate. In line with the literature^3–8,30^, all grammatical conditions that can be characterized with a regular structural pattern elicited phrase-rate tracking, whether they contained sequential regularities (V-N, V-Adv) or not (V-Adv-Alt). These grammatical tracking effects were furthermore modulated by the syntactic properties of the phrases. Moreover, we found phrase-rate tracking in conditions with sequential lexical regularities, even in the absence of grammatical structure (N-R). In all, these findings suggest that the phrase-rate tracking effect is a general neural readout of regularity tracking, which is sensitive to the presence of different levels of linguistic representation.

### 3.1 Tracking structural regularities

In both ITPC and evoked power, we observed 1-Hz phrase-rate tracking for all three conditions that contained grammatical structure (i.e., Verb-Noun, Verb-Adverb, and Verb-Adverb-Alternation), which had a central-anterior scalp distribution in evoked power. Pairwise comparisons revealed effects of syntactic properties: verb phrases with complements were tracked more strongly than verb phrases with adjuncts (V-N > V-Adv), and verb phrases with a consistent head position were tracked more strongly than verb phrases with varying head positions (V-Adv > V-Adv-Alt).

The overall pattern of phrase-rate tracking in grammatical conditions is not predicted by existing non-syntactic accounts. For instance, they cannot be explained by prosody^53^, because there is no difference between V-N and V-Adv sequences in either explicit prosodic structure (see Figure 5B) or implicit prosodic grouping (i.e., all grammatical conditions contained repetitions of verb phrases). Neither can these findings be captured in terms of differences in transitional probabilities^34,36^, because there was no reliable relationship between the magnitude of the tracking effects and bigram transitional probabilities across conditions (see Figure 4C). Alternatively, one might attribute the effect to the fact that V-N and V-Adv, as constructions, differ in frequency (e.g., the frequencies of the combinations V-N and V-Adv might be different). However, this presumes the representational capacity for syntactic abstraction in the first place. If people can compute statistics based on grammatical category combinations, they are able to represent syntactic structure extending beyond lexical representations. Frequency-based explanations of this kind must therefore be elaborated with structural language representations.

**Figure 5.**
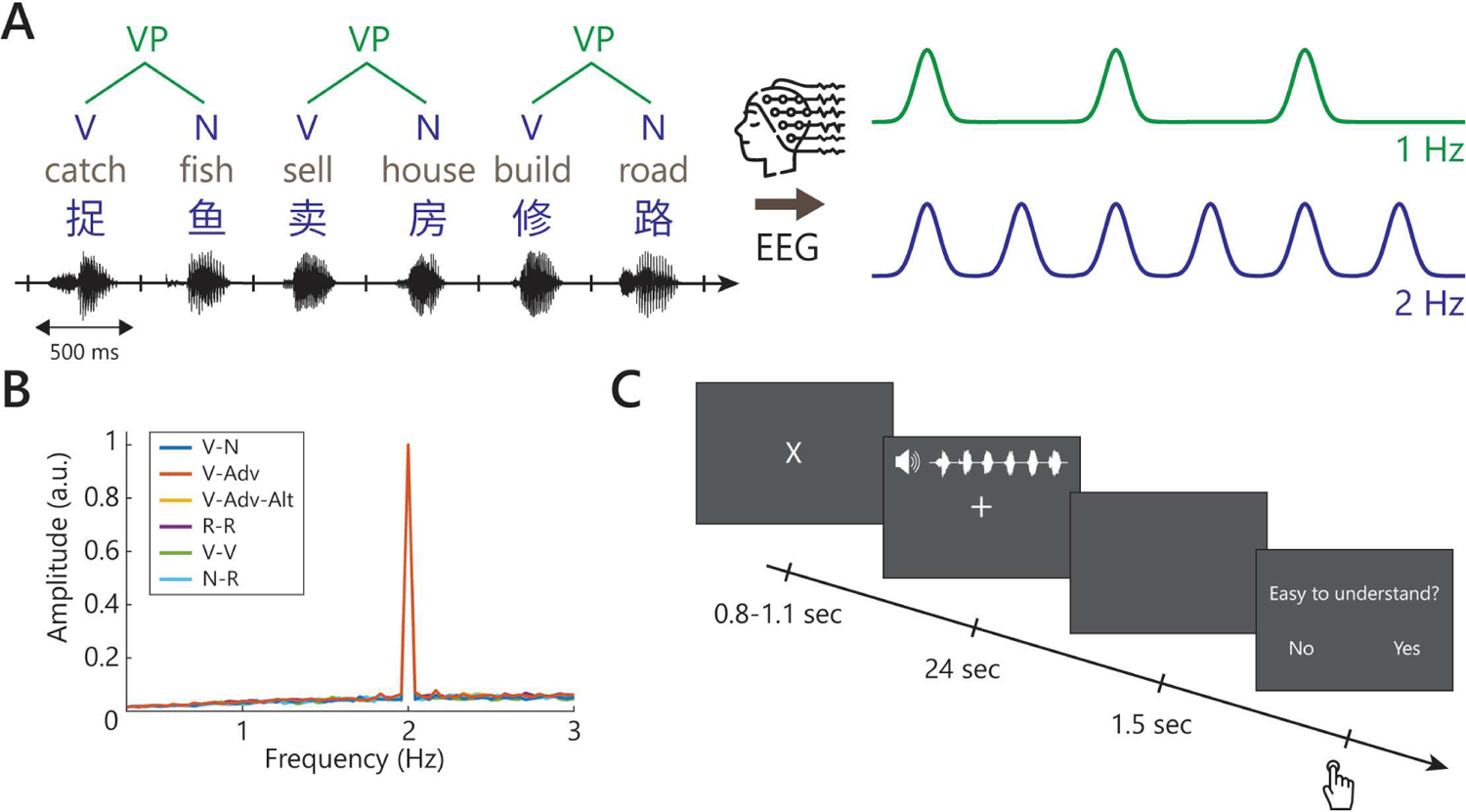
Schematic representation of the materials and experimental procedure. **(A)** Multi-level linguistic units present in connected speech sequences (left) and hypothetical neural responses to different levels of linguistic representation (right). This is a toy illustration of the temporal unfolding of monosyllabic words (both every 500ms) and phrases (every 1000ms), with hypothetical neural responses reflecting the processing of words and phrases at the corresponding rates of presentation. **(B)** The spectrum of stimulus intensity reflects the average spectral amplitude of synthesized speech signals in the frequency domain. A spectral peak was present in all conditions only at 2 Hz, corresponding to the word rate. The six conditions were not acoustically different. **(C)** Illustration of an experimental trial in one condition. (This figure includes icons from Flaticon.com)

Verb-noun phrases were tracked more closely than verb-adverb phrases. This effect of attachment type is consistent both with the theoretical distinction between complements and adjuncts and with their different processing characteristics. According to certain linguistic theories, complements and adjuncts are different in both syntax^38,39^ and semantics^37^. A potential processing-based explanation for increased phrase-rate tracking of complement phrases is that complements satisfy the selectional restriction of the verb, saturate its argument structure, and therefore reduce uncertainty in processing. Adjuncts, by contrast, only modify the verb without fulfilling any requirement, which does not significantly reduce uncertainty. The resulting uncertainty around adjunct attachment might yield increased variance in a stream of V-Adv structures, reducing phase coherence and hence phrase-rate tracking.

Consistently head-initial phrases were tracked more closely than sequences with varying head position (effect of head position). The tracking difference between these conditions can be explained in terms of different positions of the grammatical head, which determines the syntactic status of the phrase and guides syntactic processing^40,41,44–46^. We find the head-position effect intriguing because it shows that the brain accomplishes a seemingly dichotomous task, namely tracking *abstract* linguistic structure while preserving structural *detail*. That is, it is unlikely that the head only serves as a salient landmark that repeatedly evokes a neural response or re-aligns neural oscillations^1,10^. If this were so, we should not have obtained consistent 1-Hz phrasal tracking in the V-Adv-Alt condition given the temporal fluctuation of several hundred milliseconds (i.e., phrase-initial and phrase-final heads randomly lagged by 500ms). Instead, the head position effect calls for a mechanism that resolves temporal inconsistency by inferring linguistic structure from the signal (cf.^9^). A candidate mechanism of such kind is the “time-based binding” mechanism described in a computational model of neural oscillations^54^. This model infers structure from temporally unfolding signals to generate pulses, eventually rendering a consistent readout in the frequency domain. Under this framework, the effect of head position can be explained in terms of the temporal inconsistency associated with temporal binding: the 1-Hz peak in the V-Adv-Alt condition was lower than the 1-Hz peak in the V-Adv condition, because in the former case there was more temporal jitter in the brain’s generation of a periodic endogenous pulse (reflecting the composition of a verb phrase) in the absence of reliable physical cues. This proposal is compositional in the sense that higher-level units are hierarchically composed of low-level elements and decouples from constituent-level variance (e.g., head position inconsistency). Such a compositional theory aligns with neurobiological theories of language processing that include a combinatorial operation that hierarchically combines smaller elements into larger elements^11,15,55–59^. Our findings so far suggest that the process underlying phrasal tracking is likely sensitive to order (i.e., the timepoint at which structure building is initiated differs depending on the position of the head of the phrase) and detailed syntactic information (i.e., different types of syntactic structures elicit different degrees of tracking). These findings therefore enrich the neural tracking literature by showing how the tracking of phrases in connected speech is modulated by the specific syntactic properties of these phrases (cf. ^3,5,7,8^).

### 3.2 Tracking sequential regularities

In parallel with structural tracking effects, we observed phrase-rate tracking in conditions that exhibited a sequential lexical regularity at 1 Hz. This effect was more pronounced in grammatical conditions (V-N, V-Adv), but was also found in a condition where there is no grammatical structure at all (N-R). We interpret the N-R effect as indicating a sequential process external to grammatical structure. However, as we will discuss below, the sequential nature of the N-R effect cannot reduce all grammatical tracking effects to sequential regularities.

Effects of sequential statistics are omnipresent in language processing. During sentence processing, word-level statistical information is activated in tandem with higher-level structural information to modulate neural activity^60,61^. Statistical information can also be exploited in the absence of grammatical structure, e.g., during speech segmentation^33^. Likewise, a recent frequency-tagging study^62^ reported multi-word tracking effects for sequences without any grammatical structure, when participants were instructed to perform a task tuned to specific semantic categories of the individual words (i.e., “detect two-word chunks consisting of living or non-living nouns”). Any pattern-recognition strategy operating over sequential, word-level information will yield a phrase-rate tracking readout if the presentation rate of the word-level features coincides with the frequency of phrases. In N-R sequences, participants heard a noun every other 500-ms word, hence this condition elicits a tracking effect at the 1-Hz phrase rate. This sequential strategy^13^ suffices to explain our findings in the N-R condition.

However, it seems at odds with later studies^8^ showing no phrase-rate tracking in a “reversed phrase” condition (e.g., “fur dry skin rubs” vs. “dry fur rubs skin”) that, like our N-R condition, contained lexical regularities but no syntactic structure. A potential reason behind this difference is that the reversal of certain sentences affected the grammatical categories of their words. For instance, this study^8^ used derivational nouns involving a non-noun constituent and a suffix (e.g., the noun “teach-er”), of which a syllabic reversal breaks the desired part-of-speech pattern (e.g., “er-teach” is not a noun). Moreover, their sentences contained a number of disyllabic Chinese compound verbs, such as in the A-N-V-V sentence “red bird fly1 fly2”, which roughly means “the red bird flies”. Here, “fly1” and “fly2” are two distinct verbal morphemes that together form a verbal compound. As such, these sentences contain no lexical repetition at 2 Hz, so the 2-Hz tracking effect likely stems from a combination of phrasal composition (the NP “red bird”) and morphological composition (the verbal compound “fly1 fly2”). In the reversed version of this type of sentence (i.e., a N-A-V-V sequence “bird red fly2 fly1”), there is again no lexical repetition. Reversing the sentence also breaks the NP and the compound, which can explain why they did not observe a 2-Hz phrase-rate tracking effect for reversed sentences.

While being consistent with the N-R effect, the sequential strategy fails to explain the effects in other conditions. First of all, the 1-Hz effect in N-R had a posterior scalp distribution in evoked power, quite different from the central-anterior distribution of the effects in V-N and V-Adv (see Figure 3B). Though phrase-rate tracking effects converged in the frequency spectra, different processing strategies underlying tracking effects appear distinguishable in terms of topography. A spatial distinction underlying phrase-rate tracking was also discovered^62^, who found more widespread left-temporal activation associated with grammatical grouping as compared with non-grammatical, task-induced grouping. Second, a word-level tracking mechanism cannot explain the presence of phrase-rate tracking in V-Adv-Alt. Here, there is no consistent word-level regularity at the phrase rate at all, so the 1-Hz peak cannot come from the tracking of word-level features alone. Last, an exploratory analysis of the potential interaction between structural and sequential regularities showed that the magnitude of sequential tracking was boosted by adding structure (Figure 4B), suggesting that structural and sequential effects were not identical in nature and that the presence of grammatical structure increases the neural sensitivity to sequential regularity. While preliminary, this pattern suggests that structural and sequential information may go hand in hand in the service of extracting structured meaning from spoken language.

### 3.3 A regularity-based hybrid account of phrase-rate tracking

In this section, we motivate and elaborate on a hybrid account of phrase-rate tracking where regularities at multiple levels are required to explain neural patterns. This account receives justification directly from a comparison between the observed pattern of phrase-rate tracking and the predicted patterns under different theoretical hypotheses (see Figure 2). We consider a hybrid account most plausible, given a discrepancy between the current data and existing accounts that place their explanatory emphasis on singular linguistic factors. The difference between V-N and V-Adv raises a challenge for prosodic accounts^53^, because prosodic grouping was identical across conditions (see also more discussion^21^). Phrasal tracking in V-Adv-Alt contradicts lexical accounts^13,14^, since there was no reliable lexical repetition in this condition. And although the gradience within syntactic effects favors a syntactic explanation^3,5,8^, N-R does not contain grammatical structure at all, indicating that a purely syntactic account is not adequate either. To integrate our findings and the abovementioned accounts, we provide a hybrid explanation for the phrase-rate processing of connected speech sequences below.

In the end, we arrive at a potential account in which the brain tracks regularities stemming from syntactic structure and from sequential lexical information. Given that language processing is situated in a biological organ sensitive to multiple levels of information^11,15,63^, neither structural nor sequential regularities are unique to the neural tracking of language. Instead, we suggest rigorous dissection^9^ of, for example, auditory^14^, lexical^13^, and task-modulated aspects^62^ of this frequency-tagged readout. The prospect of such an idea is supported by the recent progress of neuro-computational language models in explaining neural data^19,64,65^. To summarize, this study shows that phrase-rate neural tracking is sensitive to regularities at multiple representational levels. As regularities can be computed over both the sequential pattern of lexical information and the hierarchical structure of phrasal units, our findings call for a neurobiological account of language processing where the brain leverages regularities computed over multiple levels of linguistic representation to guide rhythmic computation, and which seeks to integrate the contributions of structural and sequential information to language processing and to behavior.

## 4. Methods

### 4.1 Participants

Twenty-two native speakers of Mandarin Chinese participated in this study (15 females, mean age = 27.3, SD = 4.19) at the Max Planck Institute for Psycholinguistics (MPI). All subjects were reimbursed for their time (27 euros for 2.5 hours). Written informed consent was obtained prior to the experiment. This study was approved by the Ethics Committee of the Faculty of Social Sciences at Radboud University Nijmegen. The entire procedure was performed in accordance with relevant guidelines and regulations.

### 4.2 Stimuli and design

Participants listened to streams of monosyllabic Mandarin Chinese words that were isochronously presented at 2 Hz (see Figure 5A; the full stimulus list can be found in the Supplementary Materials). There were six conditions in total (see Table 1 for examples), consisting of repetitions of: verb-noun phrases (V-N), verb-adverb phrases (V-Adv), verb-adverb phrases with varying order (V-Adv-Alt), combinations of a noun and a pseudorandom word (N-R), pseudorandom words (R-R), and verb-verb combinations (V-V). In the N-R and R-R conditions, R words were chosen pseudorandomly rather than randomly because we had to make sure that no grammatical combinations were present. For instance, the R word in the N-R condition could never be a verb, as that could generate a subject-verb sequence. In the first three conditions (i.e., V-N, V-Adv, and V-Adv-Alt), the combination of two adjacent words yields two-word phrases, which occur at 1 Hz. These are the grammatical conditions. In the latter three (i.e., N-R, R-R, and V-V), no grammatical combinations could be formed by combining adjacent words, which makes them non-grammatical conditions. Furthermore, the conditions V-N, V-Adv and N-R contain a sequential regularity at 1 Hz, because one or more of their parts of speech are repeated every 1 second. The conditions V-Adv-Alt, R-R, and V-V contain no sequential regularities at 1 Hz.

Monosyllabic words were first synthesized using Google Text-to-Speech (Mainland Standard Chinese, WaveNet-C, male voice, speech rate = 0.75). Their duration was adjusted to exactly 500 ms by either padding zeros to the edges or truncating exceeding signals. After length normalization, a sinewave ramping window was applied to the first and last 10% of each signal to make the speech envelope more natural. Then, 48 words were concatenated into streams of 24 seconds. Figure 5B shows the power spectrum of the auditory streams of each condition, which was computed by first applying a Hilbert transform to the half-wave rectified speech signal to extract the temporal envelopes, and then applying a discrete Fourier transform to the down-sampled (200 Hz) envelope of the stimuli. The power spectrum of all conditions contains a clear peak at 2 Hz only, showing that the acoustic signals only contain information at the word rate and that the conditions cannot be distinguished acoustically.

For each of the six conditions, we created 288 unique two-word combinations. These two-word combinations formed syntactic phrases in the grammatical conditions and non-syntactic ‘phrases’ (two-word combinations) in the non-grammatical conditions. Each phrase was repeated twice in different halves of the experiment, so there were 576 phrases per condition in total. These were divided over 24 streams per condition, each of which contained 24 phrases (i.e., 48 monosyllabic words per condition). These streams were then distributed over 24 blocks, with each block containing one stream from each of the six conditions. The order of conditions within each block was pseudo-randomized, and the order of the blocks across participants was counterbalanced.

### 4.3 Procedure

Participants listened to speech streams played through loudspeakers in a sound-proof and electromagnetically shielded room. They were instructed to sit still, blink normally and pay attention to the audio. After audio playback, they had to indicate by button press whether they thought the previous stream was “easy to understand”. Participants were implicitly informed through feedback in a practice block that grammatical sequences are easier to understand than non-grammatical sequences. This behavioral task served to keep participants’ attention focused on the stimuli in a natural way without invoking artificial and task-specific grouping strategies. We chose not to use a phrase- or syllable-detection/monitoring task^3,4^ after observing in pilot runs that such tasks make participants inclined to strategically group every two or four syllables, even in non-grammatical conditions. The experiment was divided into 24 blocks. Participants were allowed to take short breaks after each block. The interval between trials ranged randomly from 800 to 1100 ms. The structure of each trial is illustrated in Figure 5C.

### 4.4 EEG recording

EEG signals were recorded using an MPI custom ActiCAP 64-electrode montage (Brain Products, Munich, Germany), of which 59 electrodes were mounted in the electrode cap. Horizontal eye movements were recorded by two electrodes placed on the outer canthi of the left and right eyes. Eye blinks were recorded by an electrode placed below the left eye. One electrode was placed on the right mastoid (RM), the reference electrode was placed on the left mastoid (LM) and the ground electrode was placed on the forehead. The EEG signal was amplified through BrainAmp DC amplifiers, referenced online to LM, sampled at 500 Hz and filtered with a passband of 0.016-249 Hz. The impedance of each electrode was kept below 25 kΩ by applying electrolyte gel prior to data recording.

### 4.5 EEG preprocessing

EEG signals were preprocessed and analyzed with the Fieldtrip toolbox^66^ in the Matlab environment (Mathworks Inc., version 2021b). The EEG data were filtered with a 0.3-25 Hz bandpass filter and re-referenced to the average of the left and right mastoids (LM/RM). Channels that were malfunctioning or showed excessive drifts across the entire experiment were removed and then interpolated using the weighted average of their neighboring channels. Then, the data were epoched into single trials, from the onset of the third phrase (t = 2s; to avoid transient auditory ERP responses at stimulus onset) to the end of the sequence (t = 24s). Whole trials were rejected only if there was a consistent electrode-level artifact, such as excessive muscle movements or jumps. We used independent component analysis to regress out artifacts resulting from eye blinks and eye movements. Data from two participants were rejected during preprocessing. The data of one of them was very noisy due to excessive head movements (over 60% of trials rejected). The other participant did not show reliable tracking at the syllable rate, which is at odds with robust findings that acoustic input reliably evokes spectral peaks at the corresponding frequency of presentation^3–5,8,62^. We excluded this participant because we consider syllable tracking to be a prerequisite for phrase tracking.

### 4.6 Spectral analysis

The preprocessed data were converted into the frequency domain by applying a discrete Fourier transformation with a frequency resolution of 1/22 Hz (after removing the first two phrases, the duration of each trial was 22 seconds). We computed evoked power and ITPC between 0.3 and 5 Hz. Evoked power was computed for each condition by first averaging over all trials in each condition and then applying a Fourier transform. ITPC at frequency *f* was computed as the mean of the squared sum of Fourier coefficients at *f*, which can be calculated with the formula:

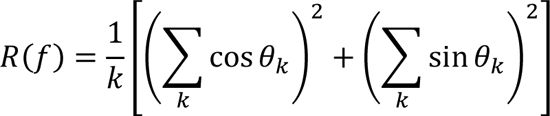

where *k* denotes the number of trials for each condition and θ_k_ is the phase angle of the complex-valued Fourier coefficient.

### 4.7 Statistical analysis

We ran statistical tests on both power and ITPC at 1 and 2 Hz. For power, we tested if the amplitude at the target frequency was higher than the average of three neighboring frequency bins to its left. By only including left-neighboring bins, this analysis takes into account the 1/f trend in power spectra. For ITPC, we tested if the ITPC at the target frequency was higher than the average of three neighboring frequency bins on each side.

Since power and ITPC in different frequency bins were not normally distributed, we ran non-parametric random permutation tests for peak detection. In each permutation, group labels (e.g., “target”, “neighbor”) were randomly assigned to observations from a merged sample containing all groups, resulting in permuted groups. Then, an arbitrary test statistic (here, we used the difference in mean between groups, denoted as Δ) was calculated for the permutation groups. Via 10,000 permutations, we created a sampling distribution of the values of the test statistic and calculated the probability of observing the actual experimental value under this sampling distribution.

After detecting peaks in each individual condition, we first performed planned pairwise comparisons (paired samples t-tests) on ITPC between all conditions that showed a 1- or 2-Hz spectral peak to evaluate the contribution of detailed structural and/or sequential configurations. A false discovery rate (FDR) correction was performed after all pairwise comparisons to correct for multiple comparisons^67^.

## Data Availability

Supplementary materials are uploaded with submission. The raw EEG dataset generated and used in the current study is available upon request.

## Acknowledgements

We thank Fan Bai for generously sharing his analysis scripts. AEM was supported by an Independent Max Planck Research Group and a Lise Meitner Research Group “Language and Computation in Neural Systems”, by NWO Vidi grant 016.Vidi.188.029 to AEM, and by Big Question 5 (to Prof. dr. Roshan Cools & Dr. Andrea E. Martin) of the Language in Interaction Consortium funded by NWO Gravitation Grant 024.001.006 to Prof. dr. Peter Hagoort. CWC was supported by NWO 016.Vidi.188.029 to AEM.

## Author Contributions

J.Z., A.E.M., and C.W.C. conceptualized the work. J.Z. and C.W.C. designed the experiment. J.Z. carried out the experiment. J.Z. and C.W.C. performed data analysis, prepared the figures, and wrote the first draft of the manuscript. J.Z., A.E.M., and C.W.C. revised the manuscript.

